# Experimental Confirmation of Osteoarthritis Repair In Vivo Using Epigenetic Reprogramming with Small Chemical Molecules

**DOI:** 10.1101/2025.03.12.642753

**Authors:** S Muratkhodjaeva, J Muratkhodjaev, T Aripova

**Affiliations:** Institute of Immunology and Human Genomics Academy of Sciences of Uzbekistan

**Keywords:** epigenetic reprogramming, aging, small chemical molecules cocktail, osteoarthritis, postmenopause

## Abstract

Osteoarthritis (OA) is a progressive degenerative joint disease that significantly impairs mobility and quality of life, particularly in aging populations. Current therapeutic approaches primarily focus on symptom relief, but fail to address the underlying mechanisms of cartilage degeneration. In recent years, epigenetic reprogramming has emerged as a promising strategy for cellular rejuvenation, offering new perspectives for regenerative medicine.

This study investigates the potential of a selected set of small chemical molecules (SCM) to induce epigenetic reprogramming of chondrocytes and promote cartilage regeneration in an in vivo model of chemically induced OA. The experiments were conducted on aging female rats, a translational model chosen for its relevance to human age-related OA and postmenopausal cartilage deterioration. OA was induced via intra-articular administration of trypsin, and a subset of animals was further subjected to estrogen receptor blockade using clomiphene citrate to simulate postmenopausal conditions. The SCM cocktail was administered intra-articularly to evaluate its effects on cartilage repair. Chondroitin sulfate was used as a comparative treatment control.

The results demonstrated that SCM administration led to significant improvements in joint tissue integrity, including increased cartilage thickness, enhanced synthesis of mucopolysaccharides, and restoration of chondrocyte metabolic activity. Histological analysis revealed that SCM-treated groups exhibited reduced cartilage erosion, lower inflammatory marker expression, and improved nuclear-cytoplasmic index (NCI) values, suggesting enhanced chondrocyte function. Notably, the group with estrogen receptor blockade responded more favorably to SCM treatment than the non-blocked OA group, highlighting the potential interaction between hormonal status and epigenetic reprogramming.

These findings provide strong evidence for the feasibility of epigenetic modulation as a therapeutic strategy for OA. The ability of SCMs to restore cartilage structure and function suggests a novel approach to treating age-related degenerative diseases. Further research is required to elucidate the precise molecular mechanisms underlying this rejuvenation process and optimize therapeutic protocols for clinical applications.

## Introduction

Aging is a complex biological process characterized by the gradual decline of tissue and organ function. Among the various theories of aging, the epigenetic theory plays a key role, suggesting that the accumulation of changes in the epigenetic profile of cells—such as DNA methylation and histone modifications—leads to dysregulation of gene expression responsible for tissue homeostasis (Guo J. et al., 2022; López-Otín C, Blasco MA, Partridge L, Serrano M & Kroemer G., 2013; Lu Y.R, Tian X, & Sinclair D.A, 2023; Pal S, & Tyler J.K, 2016; Yang JH et al., 2022).

These changes are particularly critical in tissues with low regenerative capacity, such as articular cartilage, where epigenetic dysregulation has been implicated in the progression of osteoarthritis (OA) (Diekman B. & Loeser R., 2024; Loeser R.F., 2009). OA is the most common joint disease, significantly reducing quality of life, especially in older individuals. Its pathogenesis involves hyaline cartilage degradation, chronic inflammation, and synovitis (Croft A.P., 2019; Long H et al., 2022). One of the key risk factors for OA in women is postmenopausal estrogen deficiency, which exacerbates inflammatory and degenerative processes in cartilage, thereby accelerating disease progression (GBD 2021 Osteoarthritis Collaborators., 2023; Ushiyama T, Ueyama K, Inoue K, Ohkubo I. & Hukuda S., 1999).

Current therapeutic approaches primarily aim to relieve pain and improve joint function but do not prevent disease progression or target its underlying causes (Berenbaum F, 2024; Nuesch E et al., 2011; Wehling P, Evans C, Wehling J. & Maixner W, 2017). Consequently, strategies focused on epigenetic reprogramming of chondrocytes—the primary cells responsible for cartilage integrity— are gaining increasing attention (Ball HC, Alejo AL, Samson TK, Alejo AM & Safadi FF, 2022).

Several studies have demonstrated the potential of epigenetic rejuvenation in various cell cultures exhibiting replicative senescence through the use of small chemical molecules (SCMs) (Pereira B et al., 2024; Yang JH et al., 2023). These molecules modulate key epigenetic mechanisms, including chromatin remodeling and DNA methylation, restoring the transcriptional profile characteristic of a younger state. In particular, **in vitro** experiments have shown that epigenetic modifiers can enhance cell proliferative potential and reduce the expression of inflammatory markers (Knyazer A. et al., 2021; Nie B. et al., 2017).

The aim of this study was to develop an **in vivo** model of induced osteoarthritis in rats to allow for the rapid and effective assessment of cartilage tissue restoration following intra-articular administration of SCMs.

When modeling osteoarthritis, it is essential to create a disease phenotype that corresponds to age-related OA without a predominant autoimmune component. To achieve this, we conducted a preliminary evaluation of chemically induced osteoarthritis in a rat model. In parallel, we simulated a postmenopausal state by administering estrogen receptor blockers. This pharmacological approach has significant advantages over surgical ovariectomy, as it minimizes disability in experimental rats, thereby reducing potential confounding effects on the study outcomes.

Estrogen receptors mediate the effects of these hormones on the function of various tissues in living organisms (Corciulo C. et al., 2021; Ge Yuxiang et al., 2019; Horkeby K et al., 2022). As previously demonstrated (Kalashnikova S.A. & Novochadov V.V., 2009), their blockade with clomiphene citrate effectively induces postmenopausal conditions in experimental rats.

Throughout the experiment, immunobiochemical parameters of the experimental rats were monitored through regular blood serum analyses. The final conclusions regarding the ability of SCMs to restore damaged cartilage tissue were based on a comprehensive histochemical analysis.

## Results

To study the effect of small chemical molecules (SCMs) on joint tissues in OA, a series of experiments were conducted to obtain reliable intermediate data in stages. For this purpose, the entire study was divided into two stages: Stage I—modeling OA in rats, and Stage II—studying the effect of SCMs on cartilage tissue in an aging organism with OA and postmenopause (Fig. 1).

**Figure 1.**
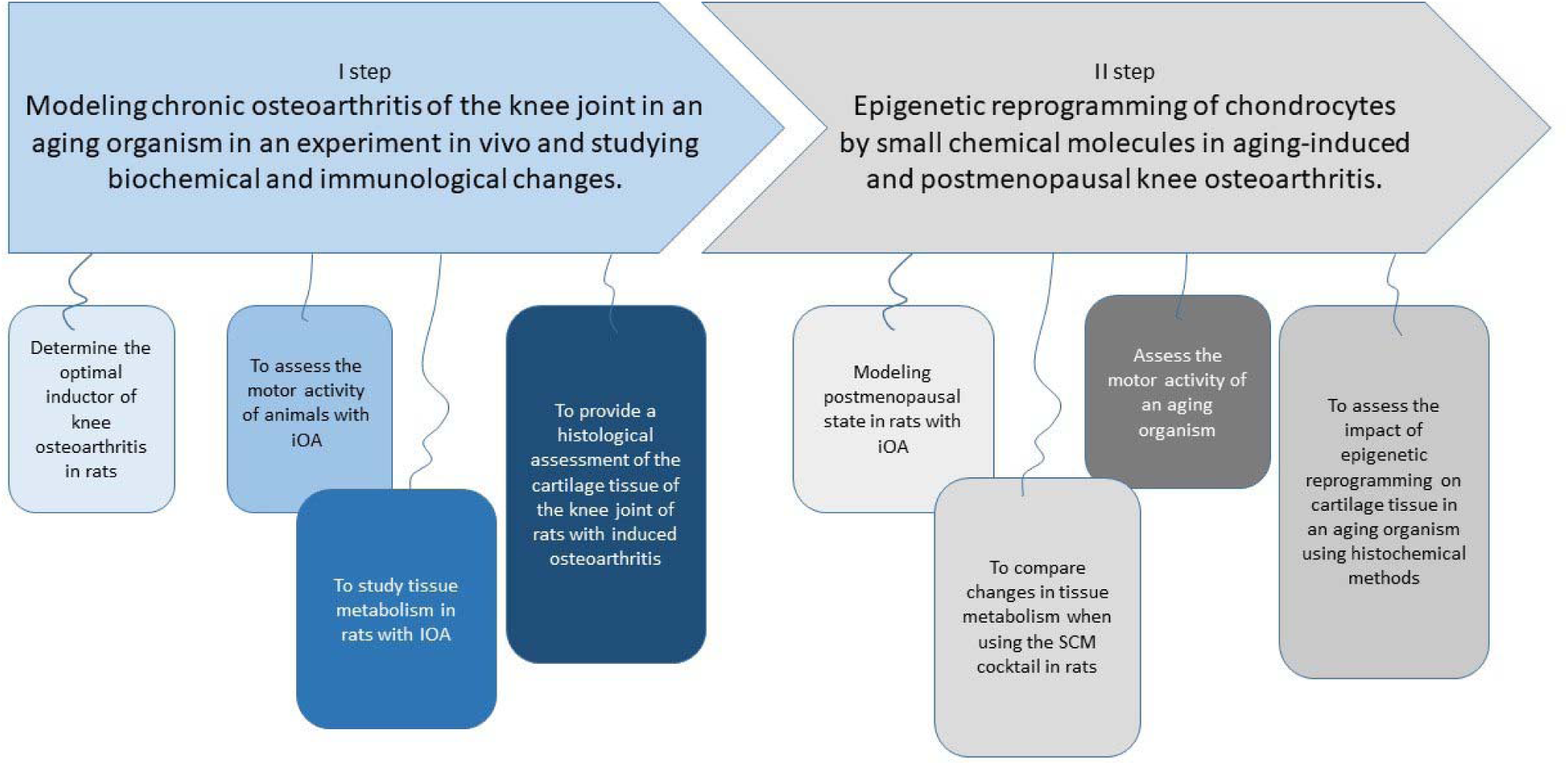
Stages of research work.

To select the optimal inducer of knee joint OA in rats, 34 female rats were randomly divided into four groups: three groups of 8 rats and one control group consisting of 10 animals. The first group of animals was injected with 0.5 ml of a 10% talc suspension, the second group with 0.1 ml of 0.1% trypsin, and the third with 0.1 ml of 1% papain. All injections were made once inside the knee joint under sterile conditions. The animals in the control group, for comparative analysis of the stress experienced, received an intra-articular injection of 0.5 ml of sterile saline. The design of the experiment is shown in Fig. 2.

**Figure 2.**
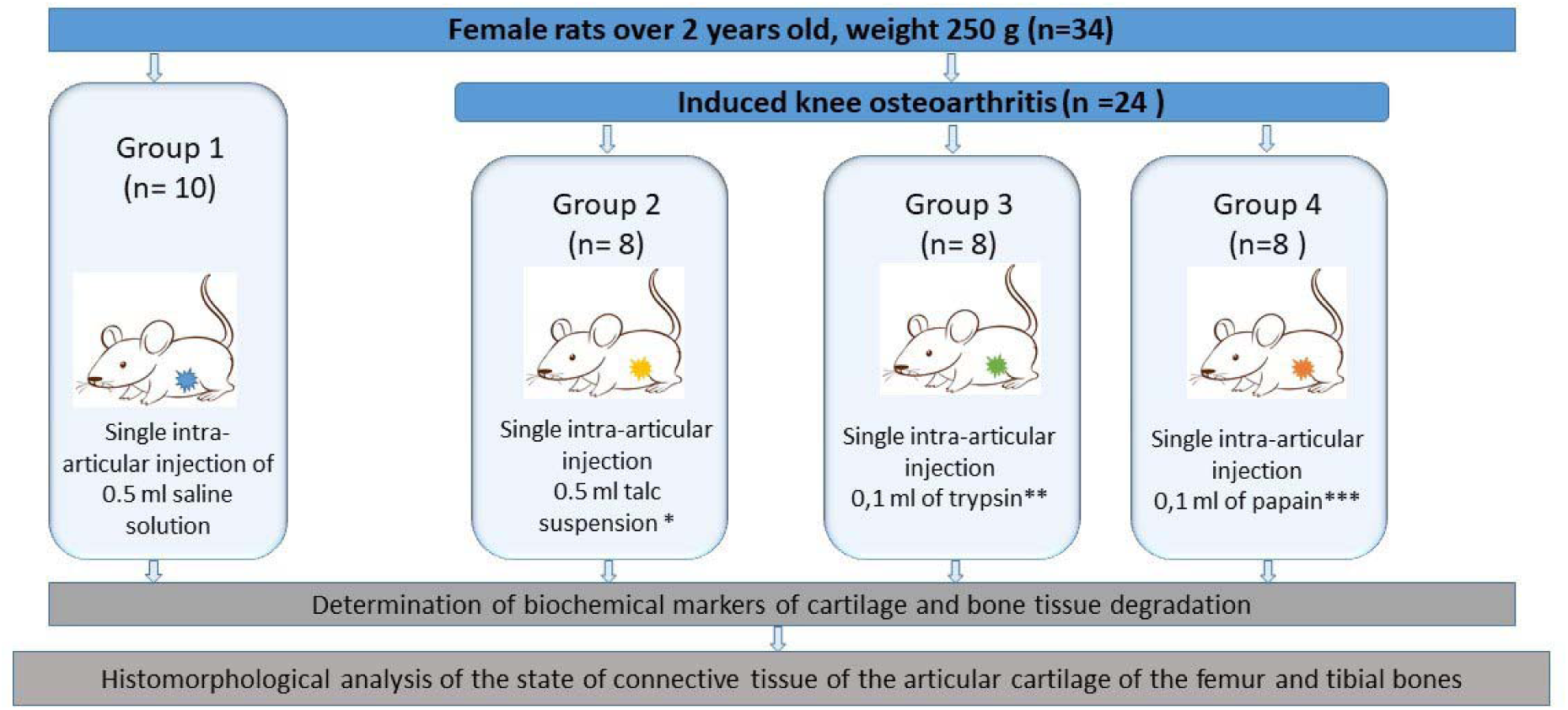
Design of experiments to verify the model of induced osteoarthritis in rats (Step I).

For 60 days after the introduction of OA inducers, the clinical condition of the animals was monitored, including animal weight, behavioral reactions (activity, mobility, lameness, gait), functional tests ("open field" test, “sprint” test), and biochemical and immunological markers (ALT, AST, ALP, vitamin D, IL-1β, TNF-α, IL-6, IL-8, and C-reactive protein).

The first clinical manifestations of developing gonarthrosis were registered as early as the second week, in the form of reduced activity, noticeable lameness, dragging of the right hind limb, and unsteady gait. Functional and motor activity, tested using the "open field" and "sprint" tests, was most reduced in Group 3 (trypsin), where the lowest indicators of horizontal and vertical motor activity, as well as movement speed, were observed.

Among the biochemical parameters, an increase in serum calcium and phosphorus levels was recorded in rats in all groups with induced osteoarthrosis compared to the control. Additionally, compared to the control group, the level of alkaline phosphatase was increased in the papain and trypsin groups (Fig. 3) but not in the talc group, which is most likely due to the effect of aluminum, a component of talc. The concentration of vitamin D in the blood of rats remained the same across all groups. The study of C-reactive protein levels revealed a significant increase in all experimental groups, indicating an acute phase of the inflammatory process. In this study, CRP levels showed a sharp increase by day 30, reaching a plateau by the end of the experiment (Fig. 4).

**Figure 3.**
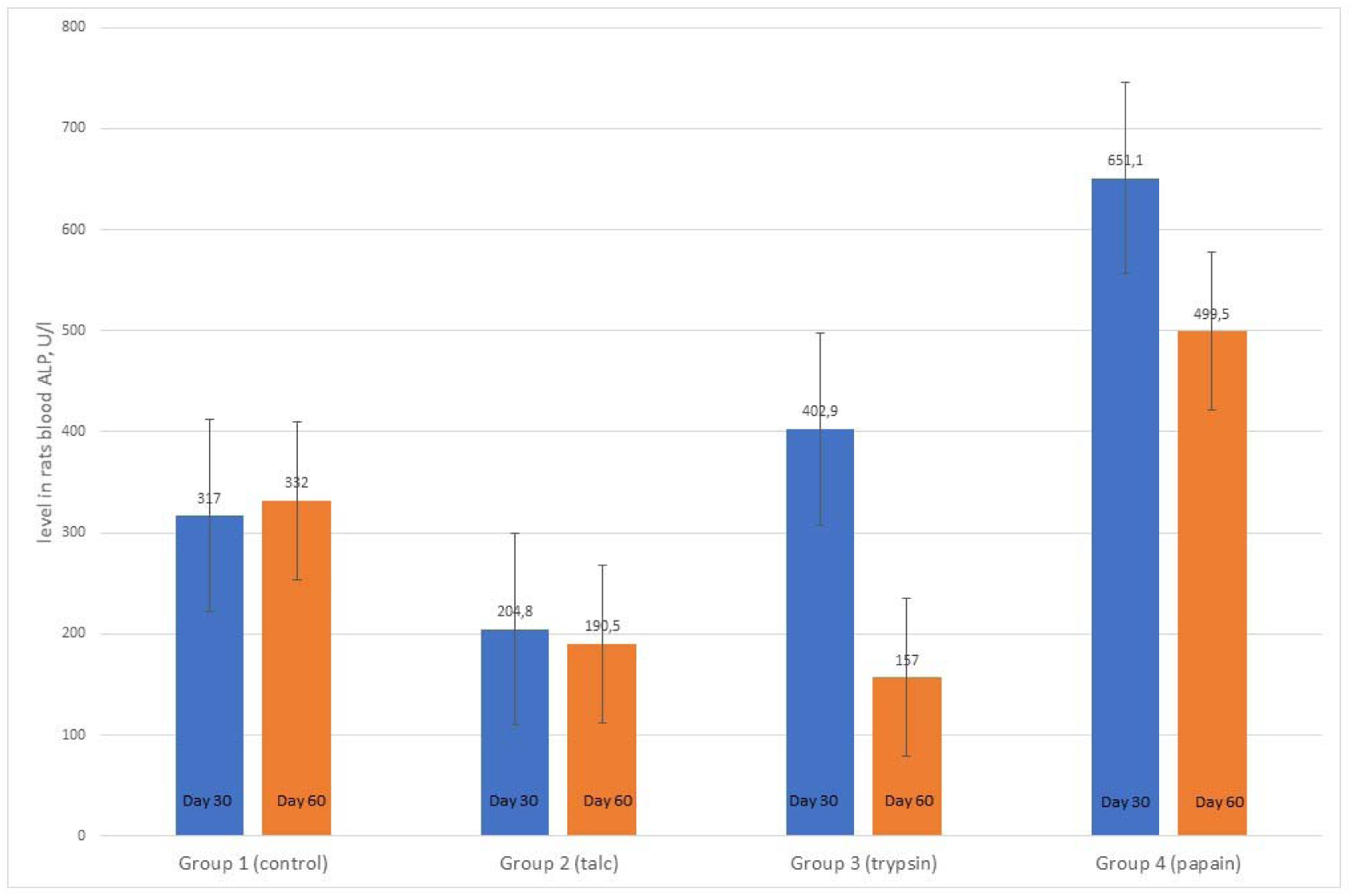
Blood alkaline phosphatase levels in rats with induced osteoarthritis on day 60 of the experiment.

**Figure 4.**
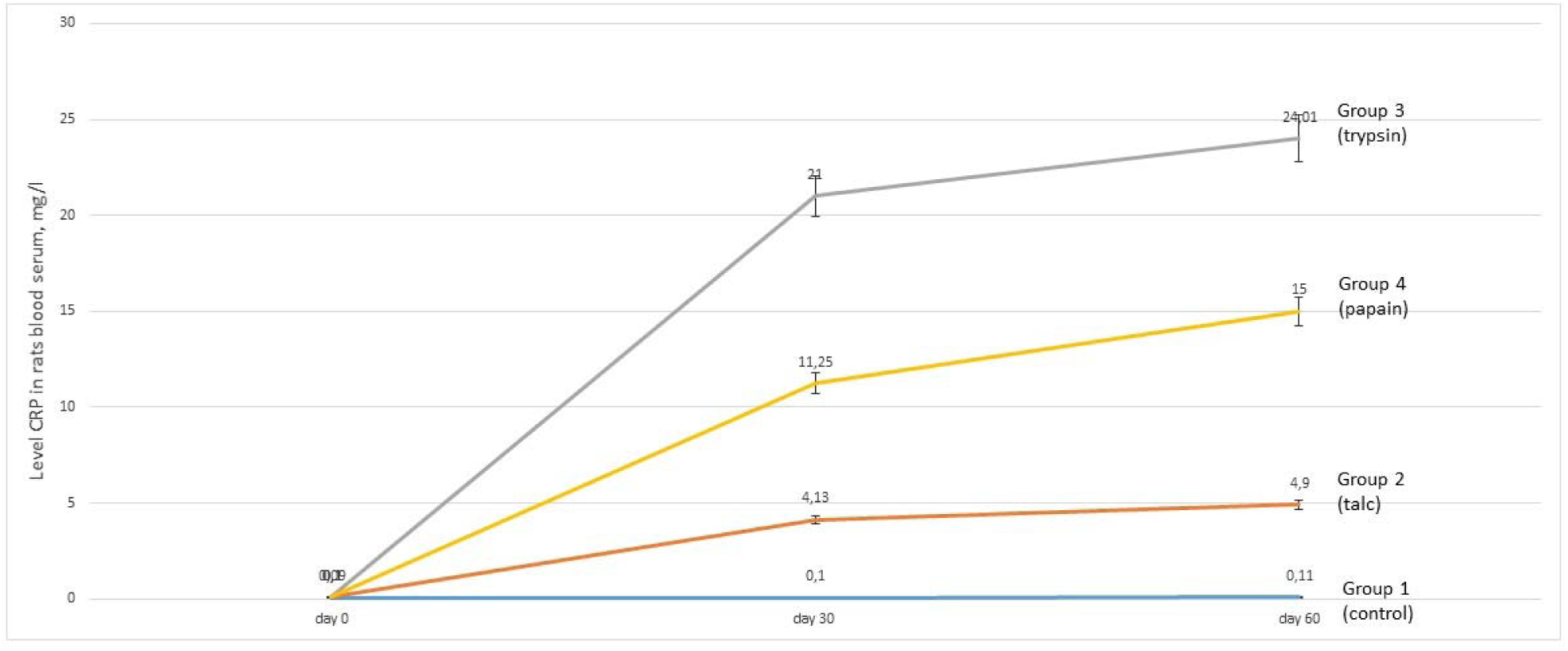
Dynamics of changes in the level of C-reactive protein in rats blood serum.

IL-6 and TNF-α levels were also elevated in all groups, peaking on day 30 of the experiment, with the highest values recorded in Group 3 (trypsin). No significant differences in IL-8 expression were observed in Group II (talc) compared to the control group; however, in Groups III and IV, a significant increase in this marker was noted. A substantial increase in IL-1β levels was detected in all groups (Fig. 5).

**Figure 5.**
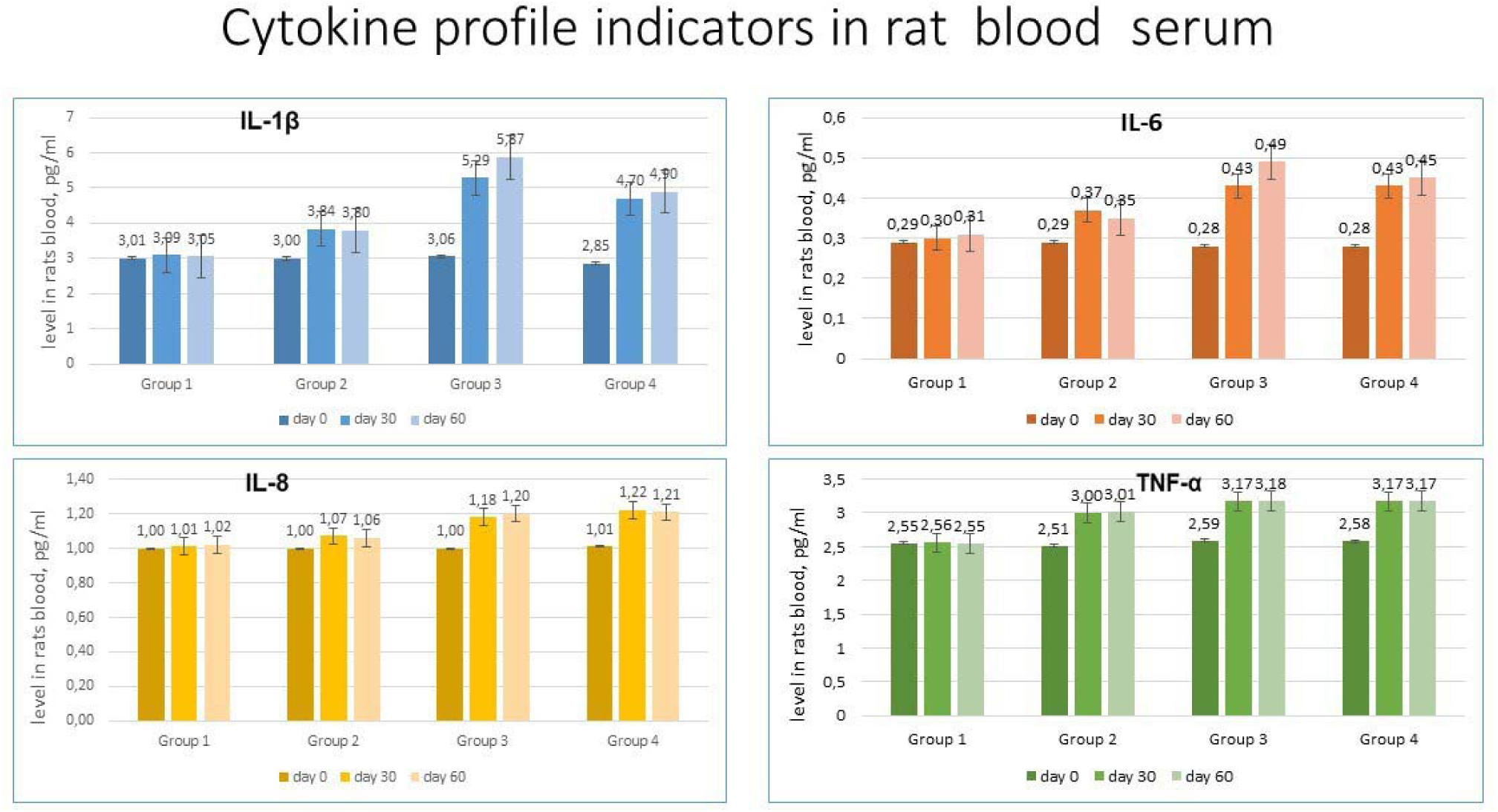
Cytokine profile indicators in rats blood serum.

Thus, the analysis of immunobiochemical parameters showed that by day 30, there was a marked increase in proinflammatory interleukin levels, particularly in the groups with enzyme-induced OA (Groups III and IV). However, by the end of the observation period (day 60), there was no exponential growth in cytokine or C-reactive protein levels, but only a slight increase. This finding suggests that a stable OA phenotype had formed by day 30 of the experiment.

After euthanasia, 34 rats were removed from the experiment for histomorphological examination of cartilage tissue. To analyze the results, histological images obtained from the same anatomical locations in different groups were compared. Comparative histological images of all study groups are presented in Table 1. In Group III, the medial, lateral, and central areas of the hyaline cartilage surface were eroded and desquamated, with pronounced chondrocyte proliferation and variations in thickness and surface roughness (Fig. 6). Pannus formation was evident by the presence of proliferating chondroblasts, around which isogenic chondrocyte groups exhibited uneven vacuolar degeneration. There was also a reduction in the homogeneity of the intercellular matrix and the detection of erosive-destructive changes extending to a depth of 1.025 mm. The joint cavity contained desquamated synoviocytes on the surface of the joint capsule and clusters of freely dispersed synoviocytes, accompanied by narrowing of the joint space, strongly indicating the development of reactive synovitis. Dystrophic and degenerative changes were found in the epimetaphyseal region, along with resorbed cystic spaces in the metaphyseal trabeculae. Since spontaneous OA does not naturally develop in rats (Gyarmati J, Foldes I, Kern M & Kiss I., 1987), it can be assumed that all observed joint tissue changes resulted from the introduction of OA inducers. Histological analysis revealed the most severe tissue alterations in Groups III (trypsin) and IV (papain). The presence of surface erosions, fissures, ulcerations, uneven hyaline cartilage thickness, matrix vacuolization, reduced cartilage staining intensity, disrupted collagen matrix structure, and blurred layer boundaries confirmed extensive destructive changes in hyaline cartilage, leading to dystrophic and degenerative processes and pannus formation.

**Figure 6.**
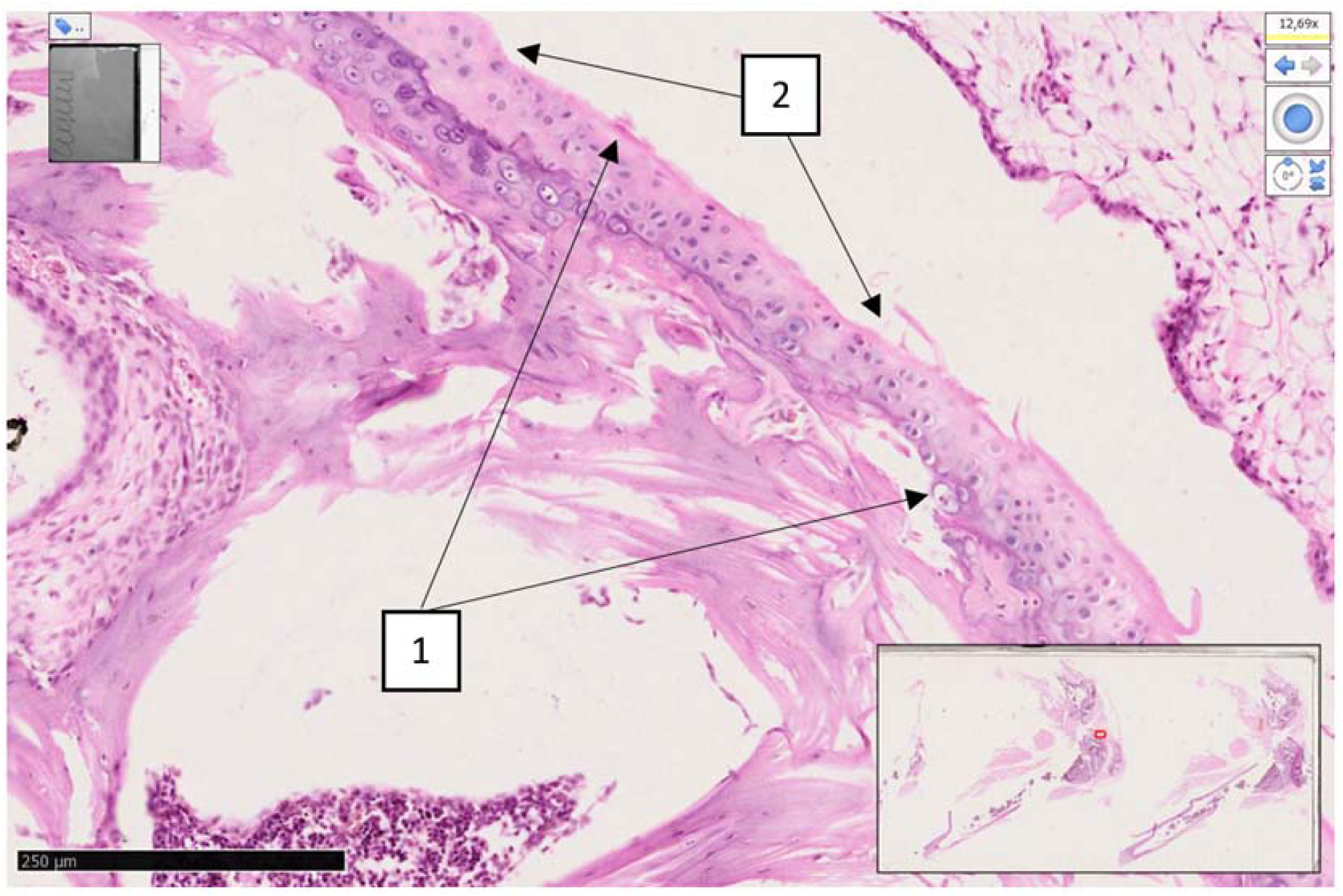
Knee joint of a rat (inducer trypsin). Sharp destructive and disregenerative changes are revealed on the lateral and medial-lateral surface of the hyaline surface of the knee joint (1), erosive-desquamative foci are revealed on the surface of the hyaline coating (2). H&E stain. Magn 10×10. The morphofunctional state of the knee joints of rats in the group with trypsin-induced osteoarthritis was determined at a level of 12 points on the Mankin scale. There are destructive and dysregenerative changes, damage to the components of the hyaline cartilage. As a result of microscopic examinations, symptoms of reactive synovitis and reactive arthritis were determined. This indicator indicates that the level of traumatism is very high, and the model used in the experimental conditions is recognized as highly effective in terms of alternative impact in this group.

**Table 1.**
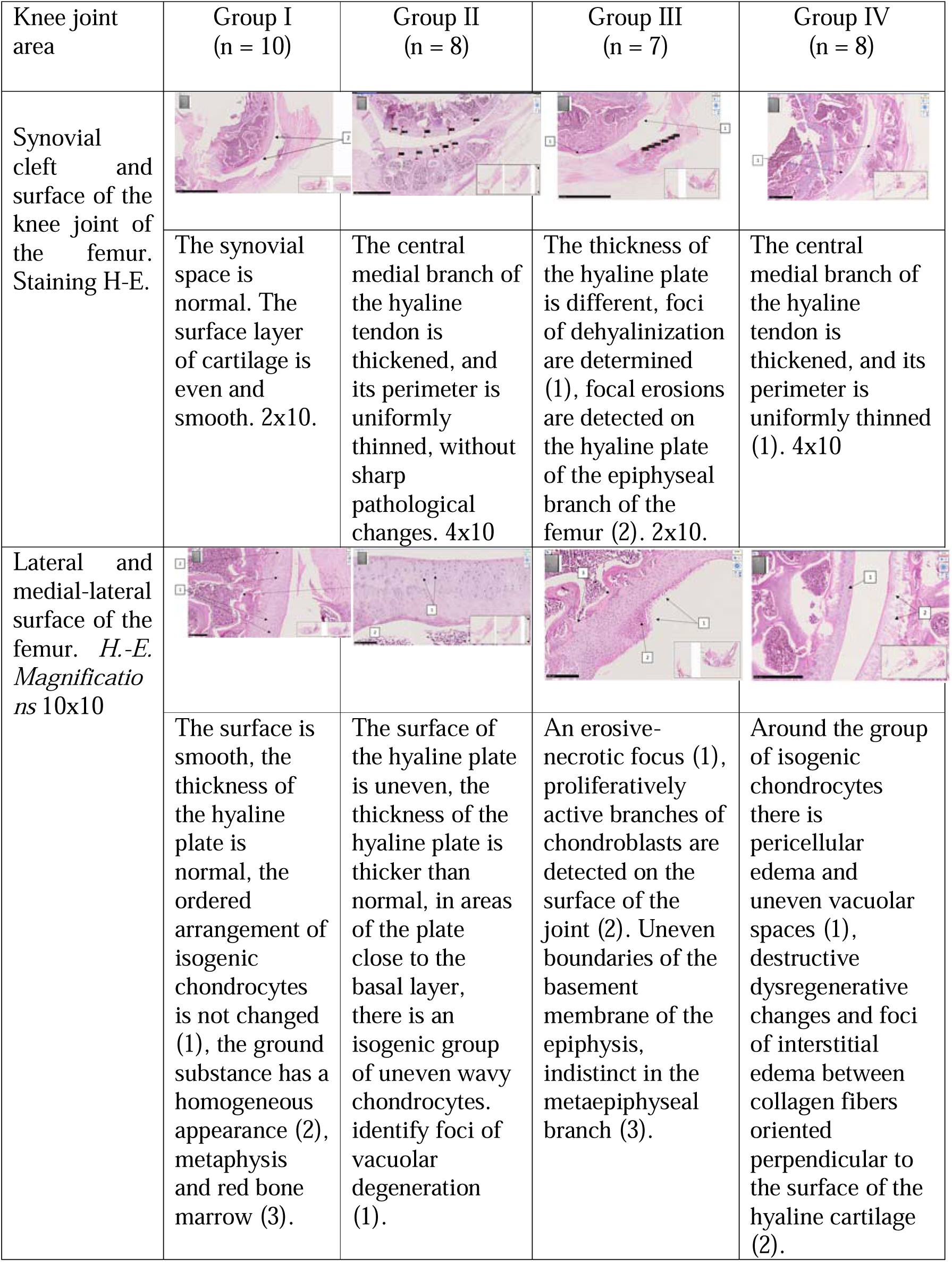
Comparative Histological Representation of Knee Joint Tissues in Rats with Chemically Induced Osteoarthritis.

| Knee joint area | Group I<br>(n = 10) | Group II<br>(n = 8) | Group III<br>(n = 7) | Group IV<br>(n = 8) |
| --- | --- | --- | --- | --- |
| Synovial cleft and surface of the knee joint of the femur. Staining H-E. | The synovial space is normal. The surface layer of cartilage is even and smooth. 2x10. | The central medial branch of the hyaline tendon is thickened, and its perimeter is uniformly thinned, without sharp pathological changes. 4x10 | The thickness of the hyaline plate is different, foci of dehyalinization are determined (1), focal erosions are detected on the hyaline plate of the epiphyseal branch of the femur (2). 2x10. | The central medial branch of the hyaline tendon is thickened, and its perimeter is uniformly thinned (1). 4x10 |
| Lateral and medial-lateral surface of the femur. H.-E. Magnifications 10x10 | The surface is smooth, the thickness of the hyaline plate is normal, the ordered arrangement of isogenic chondrocytes is not changed (1), the ground substance has a homogeneous appearance (2), metaphysis and red bone marrow (3). | The surface of the hyaline plate is uneven, the thickness of the hyaline plate is thicker than normal, in areas of the plate close to the basal layer, there is an isogenic group of uneven wavy chondrocytes. identify foci of vacuolar degeneration (1). | An erosive-necrotic focus (1), proliferatively active branches of chondroblasts are detected on the surface of the joint (2). Uneven boundaries of the basement membrane of the epiphysis, indistinct in the metaepiphyseal branch (3). | Around the group of isogenic chondrocytes there is pericellular edema and uneven vacuolar spaces (1), destructive dysregenerative changes and foci of interstitial edema between collagen fibers oriented perpendicular to the surface of the hyaline cartilage (2). |

Thus, based on the extent of joint cartilage damage observed, trypsin was selected as the optimal inducer for further studies, as it produced the most consistent OA-like pathology, closely mimicking chronic degenerative joint disease in humans.

### To study the effect of a regenerative set of small chemical molecules (SCM) on cartilage tissue in an aging organism with OA and postmenopause

At Stage II, 56 female rats meeting the same selection criteria as in Stage I were included in the study. All animals were divided into six groups of 10 animals each, including six intact rats (Fig. 7). Using the previously tested method, OA was induced with a single type of inducer—trypsin— in four groups. Additionally, a postmenopausal state was modeled in two groups by administering clomiphene in two courses of five days, leading to estrogen receptor blockade. The test SCM was administered to three groups of aging animals to evaluate its effects on healthy cartilage (Group 2), OA-affected cartilage (Group 6), and OA-affected cartilage under estrogen receptor blockade (Group 4), totaling 30 animals out of 56. Chondroitin sulfate was chosen as the therapeutic standard for OA treatment, with intra-articular injections administered at a dose of 25 mg per 250 g of body weight.

**Figure 7.**
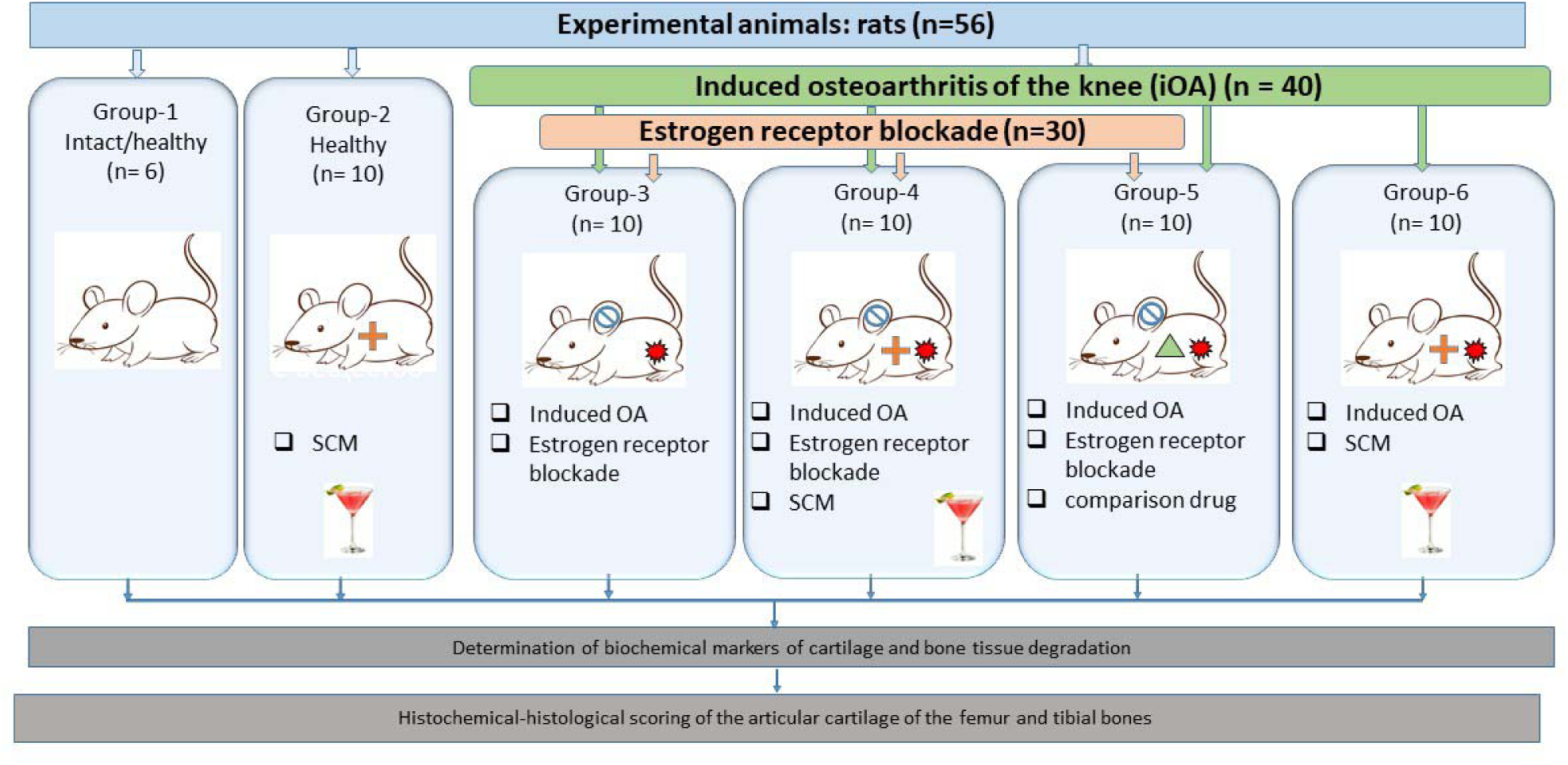
Design of studies to detect effects of a regenerative set of small chemical molecules on cartilage tissue in an aging organism with OA and postmenopause (Step II).

Throughout the 60-day experiment, a detailed observation diary was maintained, monitoring the animals’ body weight, motor activity, and periodic blood sampling for biochemical and immunological tests (see supplementary materials).

### Dynamics of Clinical and Biochemical Parameters

#### Body Weight

The weight of the rats remained stable in the intact control group and in the group of healthy animals receiving SCM (Fig. 8). However, weight loss was observed in the groups with induced OA.

**Figure 8.**
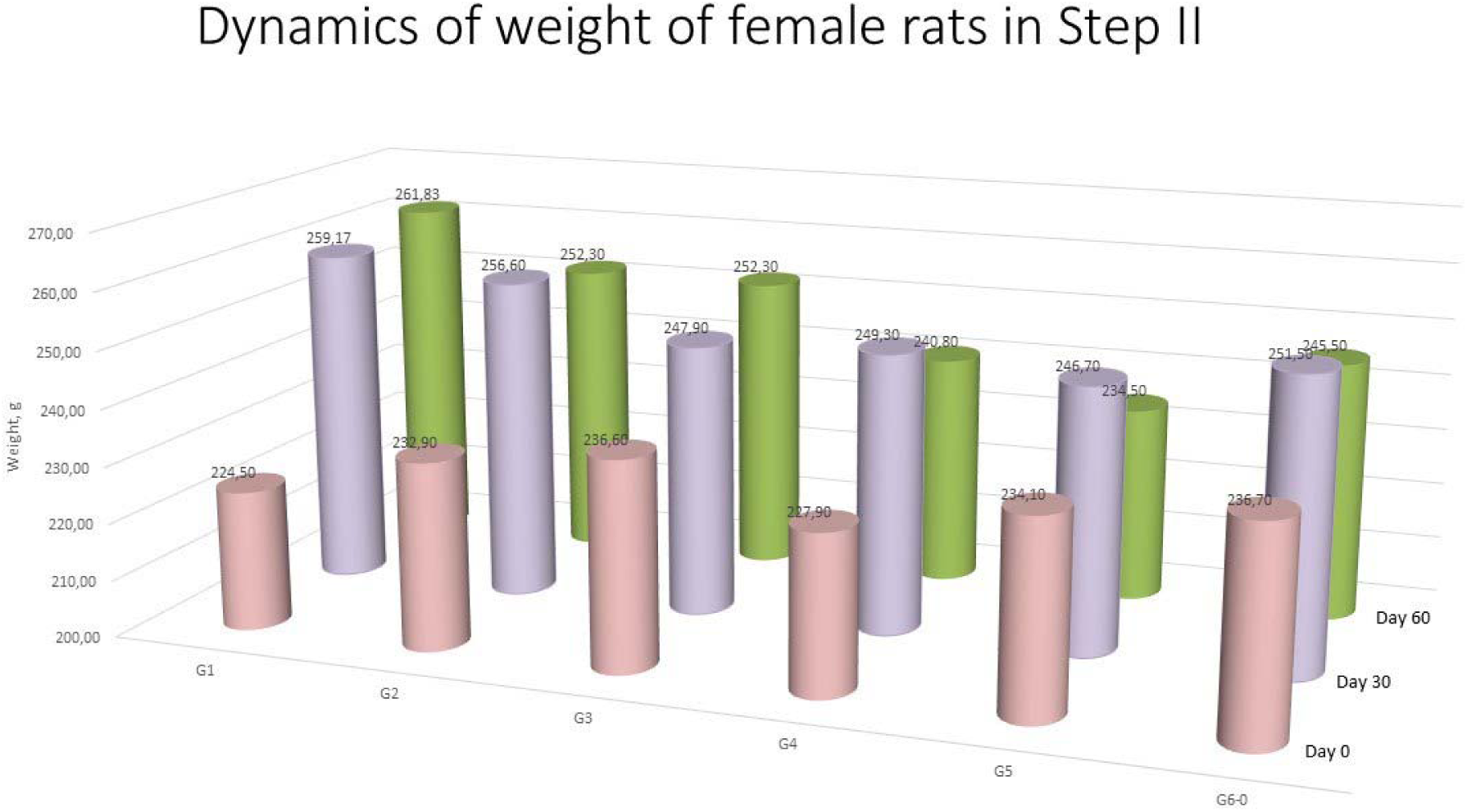
Weight of rats during experiments

#### Biochemical Parameters

Trypsin-induced OA significantly affected serum biochemical markers, including ALP, ALT, and AST. A gradual increase in ALP was observed over time in the control group (Group 1), confirming age-related changes. However, in the groups receiving SCM (Groups 2, 4, 6), ALP levels were significantly lower, indicating a slowdown in the aging process and OA progression. A reduction in ALP levels was also noted in the OA group with estrogen receptor blockade (Group 3) (Fig. 9).

**Figure 9.**
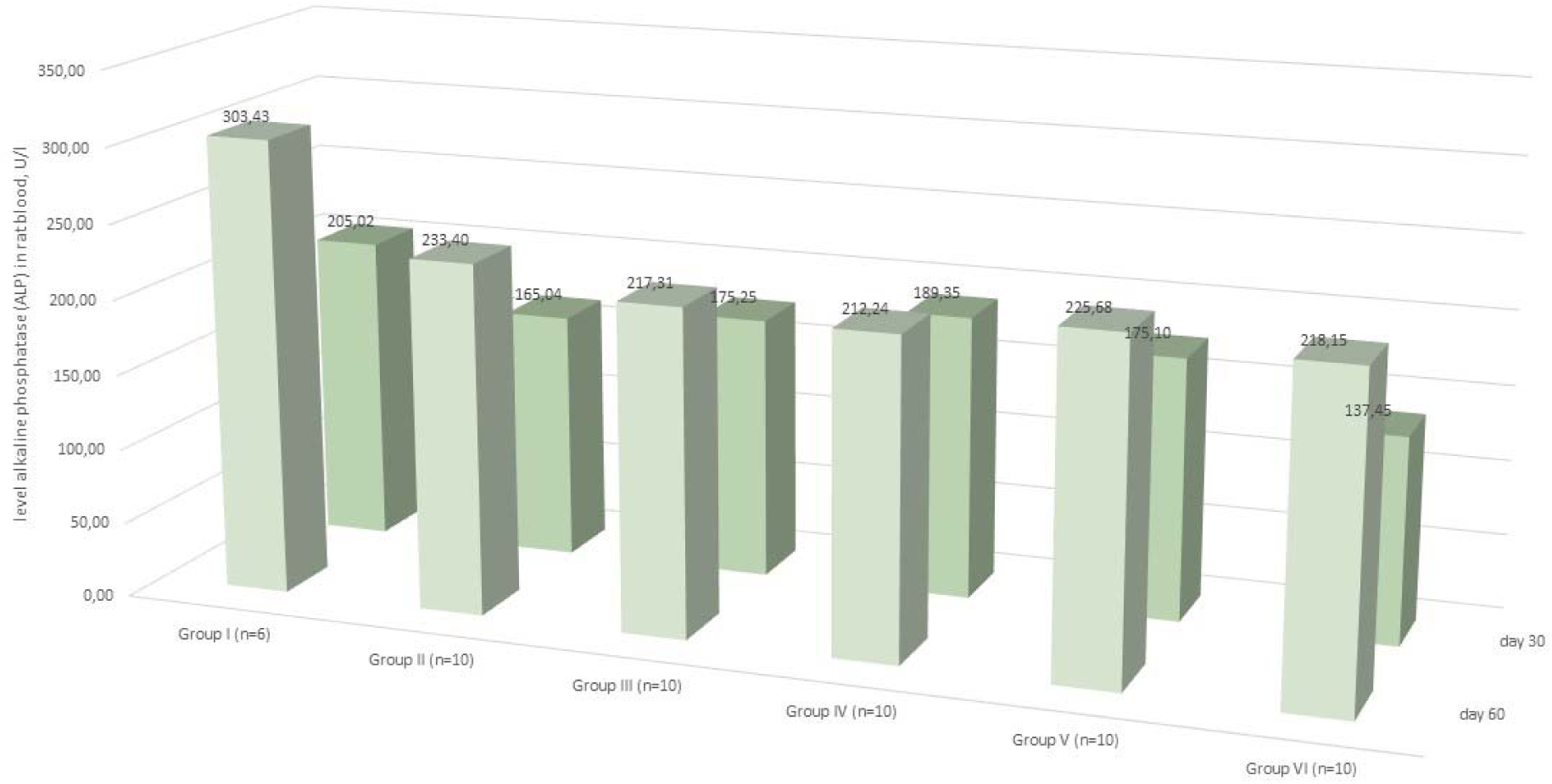
Dynamics of alkaline phosphatase levels in rats blood serum.

Maximum ALT and AST levels were recorded on day 60 in Group III, reaching 92.98 U/L and 366.15 U/L, respectively. The increase in these enzymes indicates the activation of inflammatory processes and a secretory phenotype of senescent cells, which is accompanied by the hypersecretion of cytokines and proteases. An increase in ALT levels, which indicates cell damage and inflammation, was accompanied by a decrease in ALP compared to the control group, suggesting disrupted bone metabolism (Fig. 10). A correlation was observed between alkaline phosphatase (ALP) and alanine aminotransferase (ALT) levels, given their roles in bone metabolism (Fig. 11). Notably, the group receiving chondroitin sulfate did not demonstrate significant improvements in biochemical parameters. This highlights the need for new therapeutic approaches, particularly for OA treatment in postmenopausal conditions.

**Figure 10.**
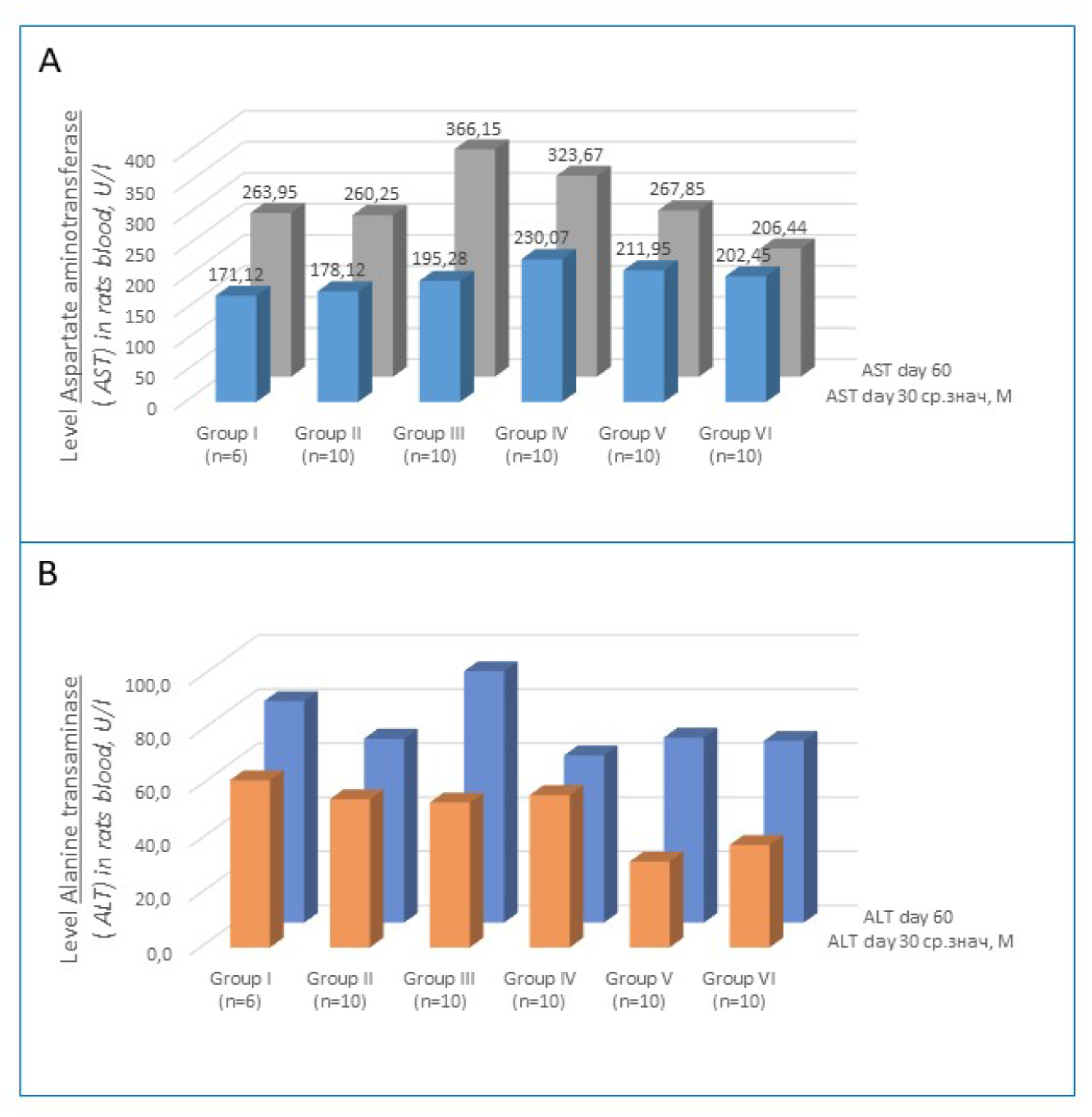
Dynamics of changes in enzymes in rats blood serum. A. AST level. 2. ALT level.

**Figure 11.**
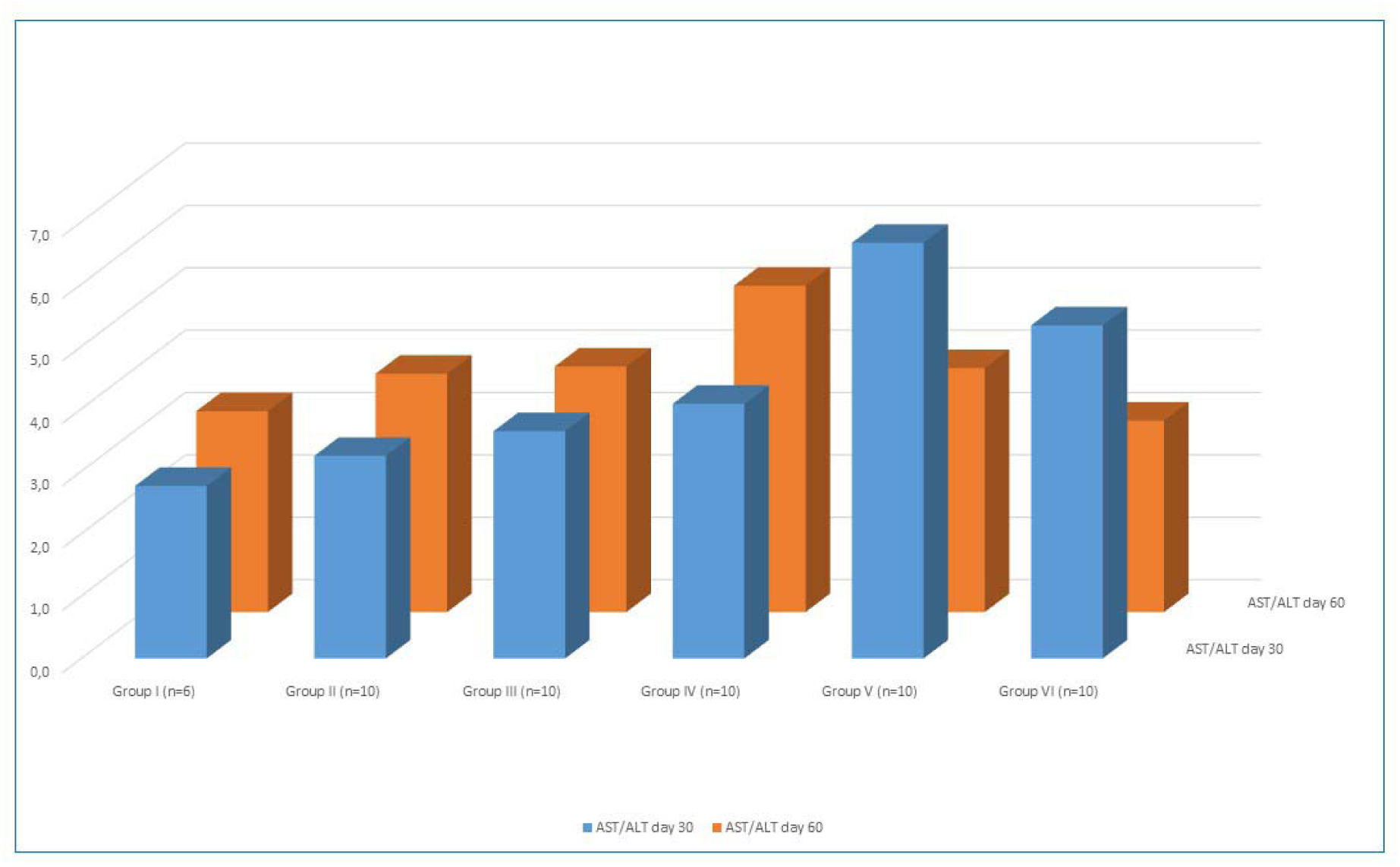
Ratio of AST and ALT levels.

### Immunological Parameters

#### Cytokine Profile

The levels of pro-inflammatory cytokines IL-1β, IL-6, TNF-α, IL-8, and C-reactive protein (CRP) reflected the development and progression of OA (see Fig. 12).

**Figure 12.**
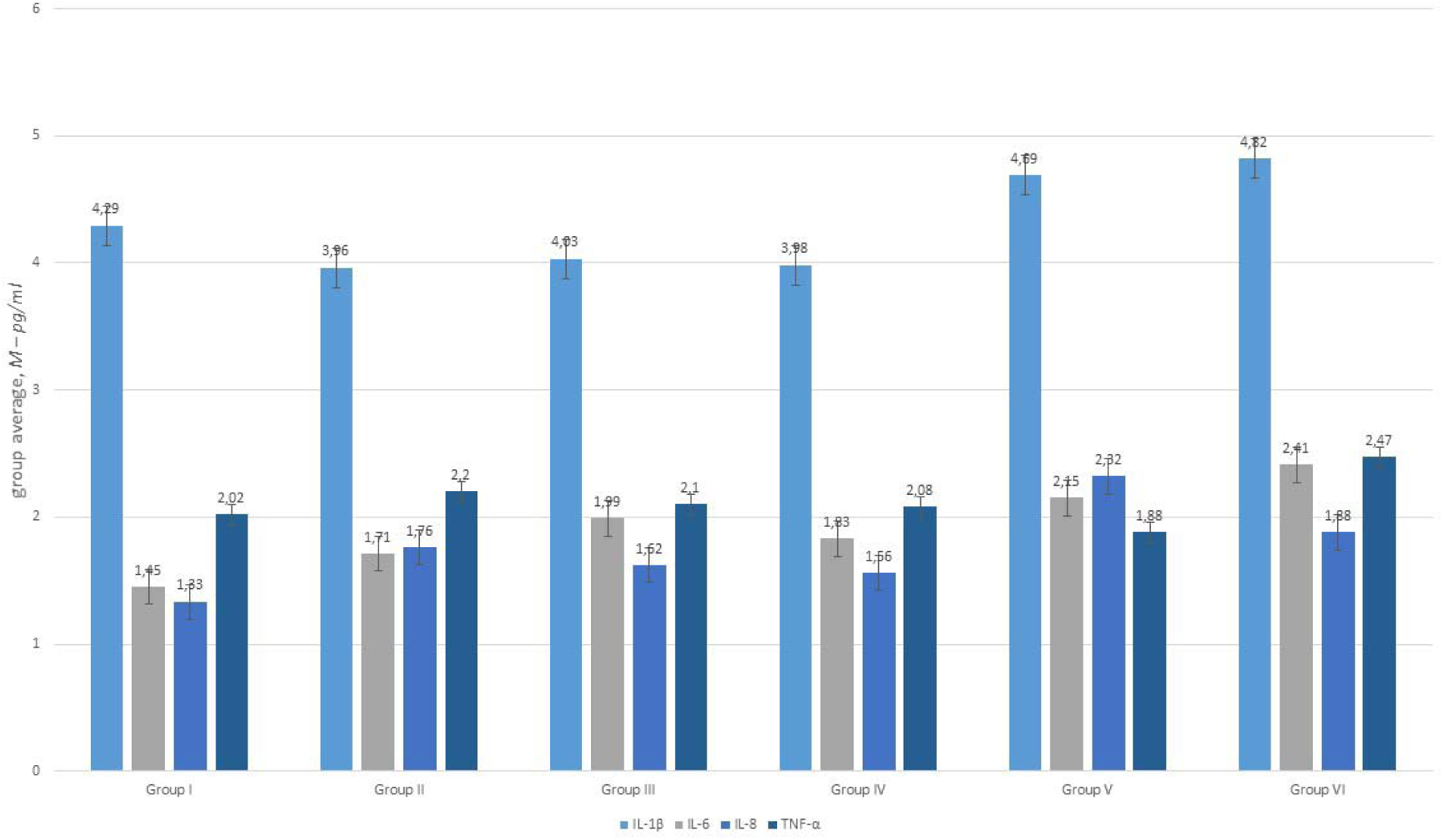
Cytokine levels in rats blood serum

The cytokine dynamics revealed:

- IL-1β peaked on day 30, coinciding with cartilage tissue degradation.
- IL-6 also peaked on day 30, particularly in the trypsin group.
- IL-8 levels were elevated from week 4 of the experiment.
- TNF-α remained consistently high in all groups, exhibiting immunomodulatory effects.

### Histological Assessment of Cartilage Tissue in an Aging Organism with Induced Osteoarthritis and the Effects of Small Chemical Molecules

Various staining methods were employed to assess cartilage integrity, including quantitative evaluation of chondrocytes and nuclear-cytoplasmic ratios. The obtained data were analyzed using the international Mankin scoring system and statistically processed. A comprehensive assessment of joint condition included key markers of chronic osteoarthritis, such as cartilage degeneration, pannus formation, joint space narrowing, synovial membrane alterations, bone tissue changes, and inflammation severity (Fig. 13).

**Figure 13.**
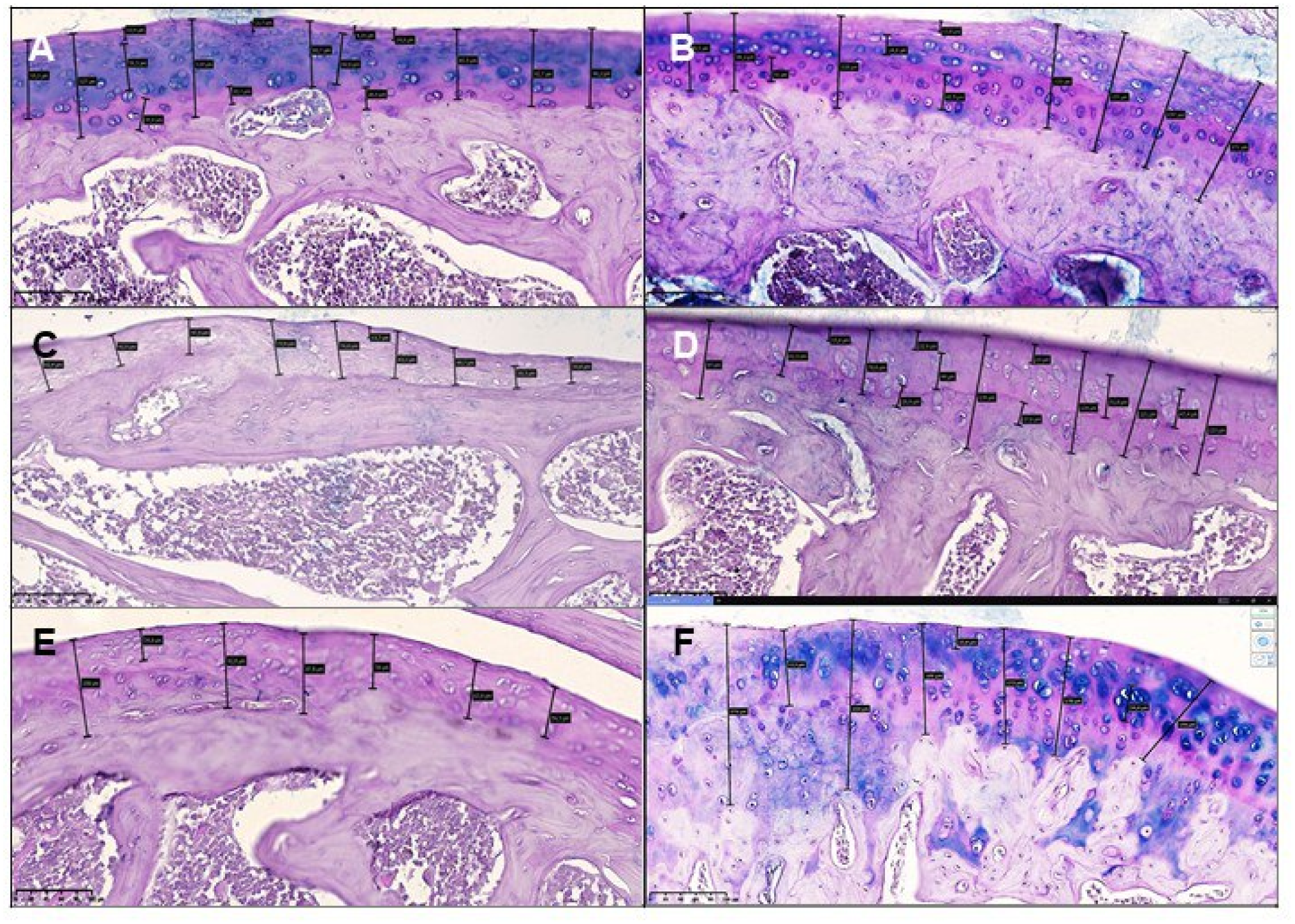
Histological morphometric data. А. Group I (intact). Epimetaphyseal region of the rat knee joint. The thickness of the hyaline plate is uniform, the cartilage zones are clearly visible, with the corresponding arrangement of chondrocytes of varying degrees of differentiation, with a predominance of young forms. H&E stain. Magn 10×10. Mankin scale score = 1. **B.** Group II (healthy + SCM). Epimetaphyseal region of the rat knee joint. Isogenic chondrocytes are located in large numbers along one trajectory in a layer of equal thickness. Mucopolysaccharides surrounding the chondrocyte cluster are stained dark blue. H&E stain. Magn 10×10. Mankin scale score = 2. **C.** Group III (iOA + postmenopause). Epimetaphyseal zone of the rat knee joint. A resorption cystic cavity is formed in the material substrate, filled with adipose tissue. The epimetaphyseal zone is invaginated into the subepiphyseal region. The bone trabeculae are small and have an atrophic appearance. Osteoclasts appear functionally active. H&E stain. Magn 20×10. Mankin scale score = 12. **D.** Group 4 (iOA + postmenopause + SCM). The surface of the rat knee joint is relatively smooth, chondrocytes in a state of proliferation and hypertrophy are found in all layers of cartilage. Chondrocytes with vacuolar dystrophy are present. Isogenic chondrocytes are grouped and located in a relatively orderly manner. H&E stain. Magn 20×10.Mankin scale score = 4. **E.** Group 5 (iOA + postmenopause + comparison drug). Epimetaphyseal zone of the rat knee joint. The border of isogenic chondrocytes varies from the metaphyseal branch to the superficial layer of the bone. The tangential surface is smooth. Reduction of Schiff-positive structures, the number of intermediates is reduced. PAS. Magn 10×10. Mankin score = 7. **F.** Group 6 (iOA + SCM). Epimetaphyseal zone of the rat knee joint. Isogenic chondrocytes are located in large numbers along one trajectory of equal thickness. There is a large amount of mucopolysaccharides around the chondrocyte clusters. H&E stain. Magn 10×10. Mankin scale score = 6.

For comparative evaluation across groups, morphometric analysis and the Mankin scale were applied (Mankin HJ, Dorfman H, Lippiello L & Zarins A., 1971; Moody HR, Heard BJ, Frank CB, Shrive NG & Oloyede AO., 2012). A quantitative assessment of cartilage and epiphyseal zones of the tibia and femur was conducted (Fig. 14). Notably, Group 6 demonstrated a significant thickening of the tibial epiphyseal zone, confirming the strong regenerative effect of SCM on OA-damaged cartilage.

**Figure 14.**
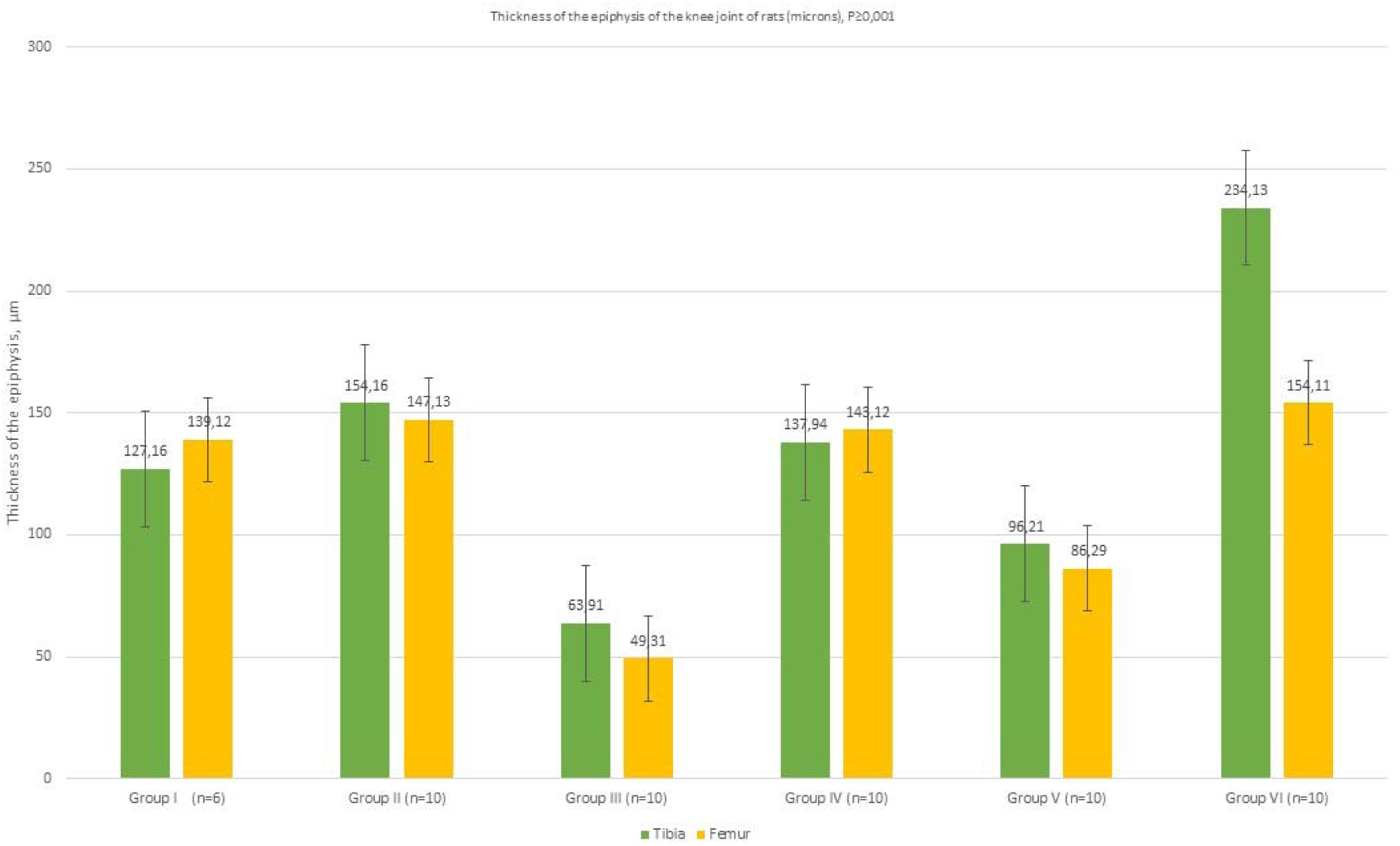
A quantitative assessment of cartilage and epiphyseal zones of the tibia and femur.

To determine chondrocyte maturation levels, the nuclear-to-cytoplasmic (N/C) ratio was calculated in various mitotically active zones of the cartilage (Fig. 15). The control group of intact animals exhibited a significantly high N/C ratio, whereas groups receiving SCM had lower N/C ratio levels than the control. This suggests only moderate activation of mature chondrocytes without the mitogenic effect of SCM components. The lowest N/C ratio values were recorded in Group III (OA + postmenopause), likely reflecting a predominance of senescent cells.

**Figure 15.**
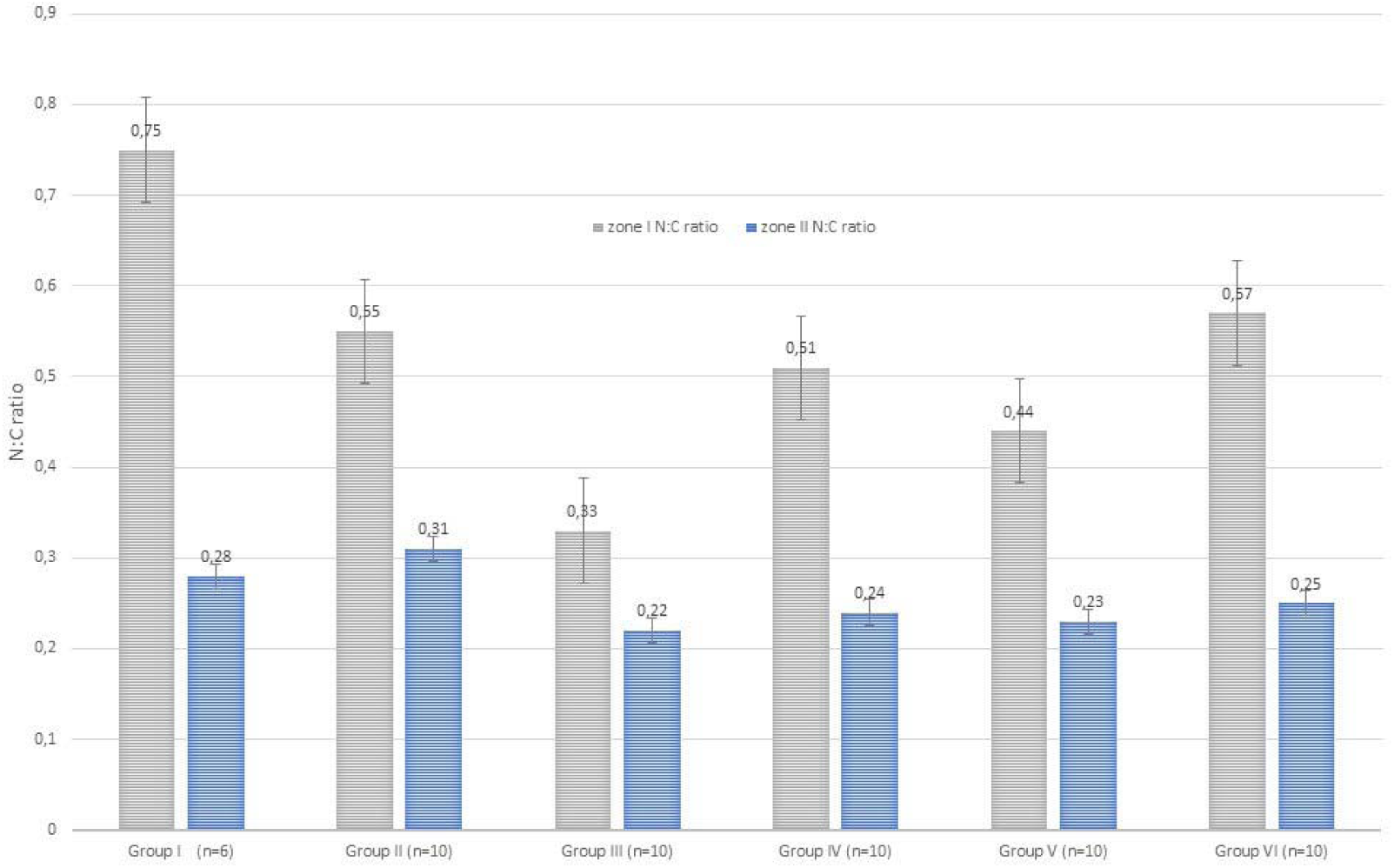
The Nuclear-Cytoplasmic (N:C) Ratio

Staining methods revealed both cartilage damage and suppressed metabolic function in the postmenopausal-simulated OA groups (Fig. 16), as well as the restoration of metabolic activity in chondrocytes following SCM administration (Fig. 17). Group III (OA + postmenopause) exhibited severe cartilage degradation, including erosions, thinning, and layer boundary loss. However, in Groups IV and VI, which received SCM, histological analysis revealed a significant improvement, indicating restored chondrocyte metabolic activity.

**Figure 16.**
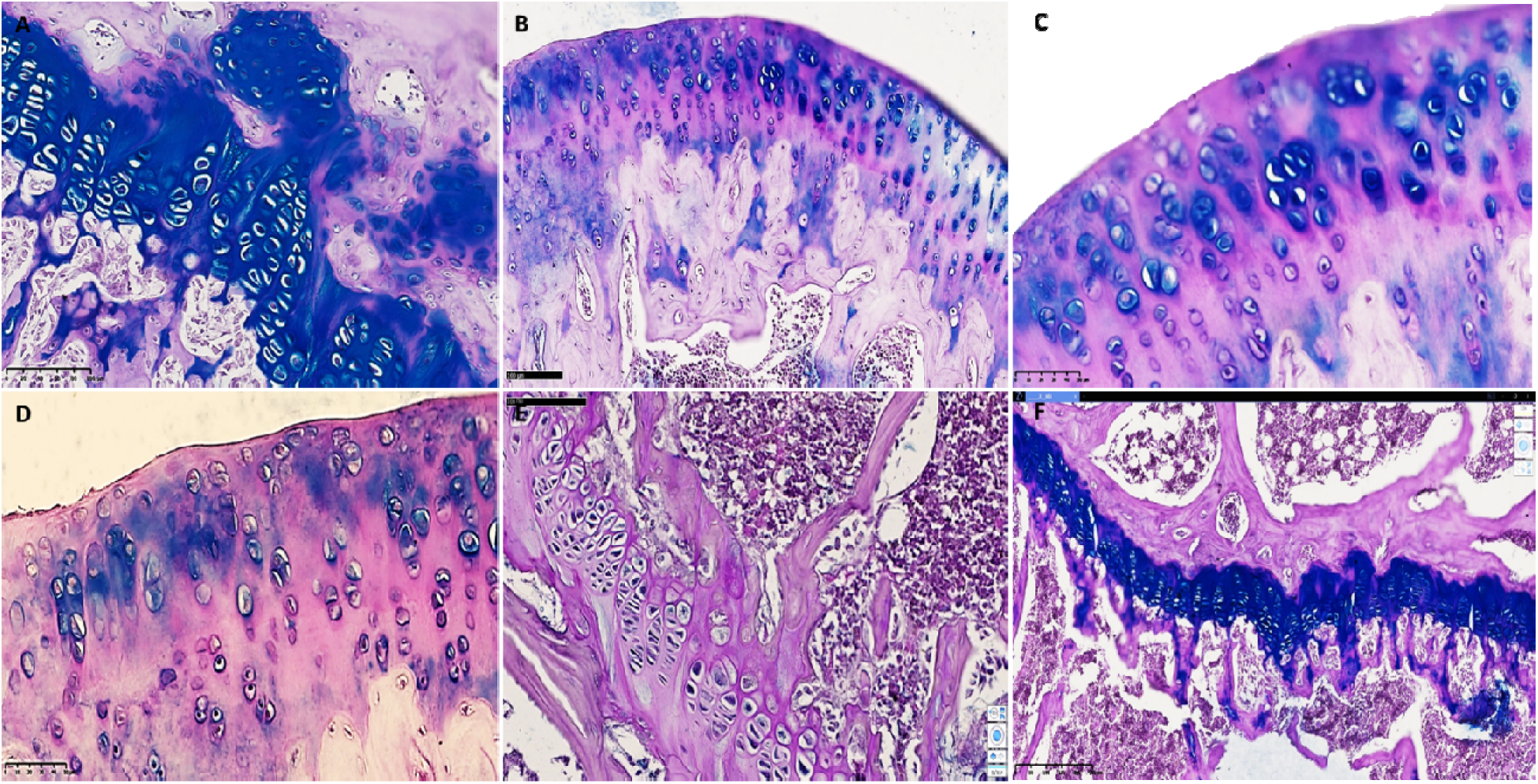
Comparison of the rat groups with knee induced osteoarthritis and estrogen receptor blockade. **A**. Group III (iOA + estrogen receptor blockade). of rats: epimetaphyseal region of the knee joint. It was found that the border of isogenic chondrocytes is shifted to the metaphyseal region, and the epimetaphyseal zone is invaginated to the subepiphyseal branch. SCHIFF-positive dark blue structures are found around chondrocytes with vacuolar dystrophy and signs of necrobiosis. Magn. 20×10. Dystrophic and disregenerative, destructive changes are revealed in bone beams. The number of osteoblasts is small, osteoclasts are increased, this process means the expansion of bone pores and the formation of resorptive cysts of enlarged foci. Cystically expanded resorption spaces indicate the development of osteoporosis processes. In the area of the joint pockets, highly developed interstitial formations with a small number of interstitial macrophages are found. **B**. Group IV (iOA + estrogen receptor blockade + SCM). The surface of the knee joint of rats of the 4th group. The surface of the joint is relatively smooth, in all rows of the hyaline cartilage cover chondrocytes in a state of proliferation and hypertrophy are found, and also, in focus were chondrocytes with vacuolar dystrophy. Isogenic chondrocytes are grouped and located relatively orderly. H&E stain. Magn. 20×10. Isogenic chondrocytes are arranged in an orderly manner, their number is 9-12 in a 200x field of view. Foci of acute proliferation of chondroblasts are revealed, cells of the isogenic group of chondrocytes have focal vacuolar dystrophy, the extracellular matrix is homogenized, the cell nuclei are stained with a basophilic dye, have a relatively uniform appearance, which indicates restored metabolic processes. **C**. Group V (iOA + estrogen receptor blockade + comparison drug). The surface of the knee joint of rats of the 5th group. The surface of the joint is relatively uneven, proliferation and hypercellular appearance are observed, multifocal vacuolar dystrophy of al rows of chondrocytes in the hyaline cartilage cover. Isogenic chondrocytes are grouped and located relatively unevenly. H&E stain Magn. 20×10.In this group, changes were observed due to multifocal destruction of the components of the joint capsule, accumulation of pyramidal lacunae of chondrocytes of different sizes around collagen fibers. On the surface of the joints, foci of erosion measuring 0.24-0.36 mm, foci of ossification in the epimetaphyseal layer of hyaline cartilage and uneven texture of the basement membrane were noted. **D**. Group III (iOA + estrogen receptor blockade). Surface of the rat knee joint. The joint surface is uneven, chondrocytes in all rows are of different sizes, with signs of vacuolar dystrophy. Isogenic chondrocytes are unevenly distributed. H&E stain Magn. 40×10. There are foci of reparative regenerat on of chondrocytes at different stages, in other areas thickenings and uneven surfaces are formed. There is very little extracellular matrix around isogenic chondrocytes, instead of it fibrous foci of coarse collagen fibers are determined. Focuses of reparative regeneration of chondrocytes at different stages are observed, in other areas thickenings and uneven surfaces are formed. In the deep layer of radial arrangement of chondrocytes, they have a hypertrophied appearance, semi-oval shape, oblong in places, are located in a disorderly manner, 3-9 groups are visible. Chondrocyte groups are subject to uneven vacuolar degeneration, the homogeneity of the interstitial substance is relatively reduced, the basophilic coloration of the cells is different, which indicates a metabolic disorder. Around chondrocytes in the state of necrobiosis and necrosis, proliferation of fibroblasts and an increase in coarse collagen fibers are observed. **E.** Group IV (iOA + estrogen receptor blockade + SCM). Rat knee joint. Epimetaphyseal region has thinned flat textured appearance, isogenic chondrocytes of the same size in the area of transition to bone trabeculae, same appearance, bone trabeculae are thickened. H&E stain. Magn. 4×10. No changes in the structure of the ligaments and tendons of the joint muscles were detected. In the epimetaphyseal zone, the isogenic chondrocytes are of the same size, the ossification boundary and the trajectory of the metaphyseal zone have the same arcuate shape, no sharp thickened zones are determined. In the epimetaphyseal and metaphyseal regions, the bone beams are of the same thickness, osteoblasts are clearly defined, their number is increased by 1.25 times, the bone beams are uniformly thickened compared to the norm, the proliferative activity of osteoblasts is increased. **F.** Group V (iOA + estrogen receptor blockade + comparison drug). Epimetaphyseal branch of the knee joint of rats of the 5th group. The border of isogenic chondrocytes varie from the metaphyseal branch to the superficial layer of the bone, multifocal vacuolar dystrophy and necrobiosis in isogenic chondrocytes are revealed around them. SCHIFF-positive method. Magn. 20×10. The structure of isogenic chondrocytes in the upper and middle sections of the hyaline layer is changed, the lacunae have a pyramidal appearance, the cells have multifocal hydropic dystrophy, and pericellular edema is detected around them. The adoption of a rounded shape by chondrocytes in this area is explained by the fact that the apoptosis process is intensified. According to histochemical tests, it was established that Schiff-positive structures are reduced, metabolism is enhanced, and the number of intermediate products is sharply reduced.

**Figure 17.**
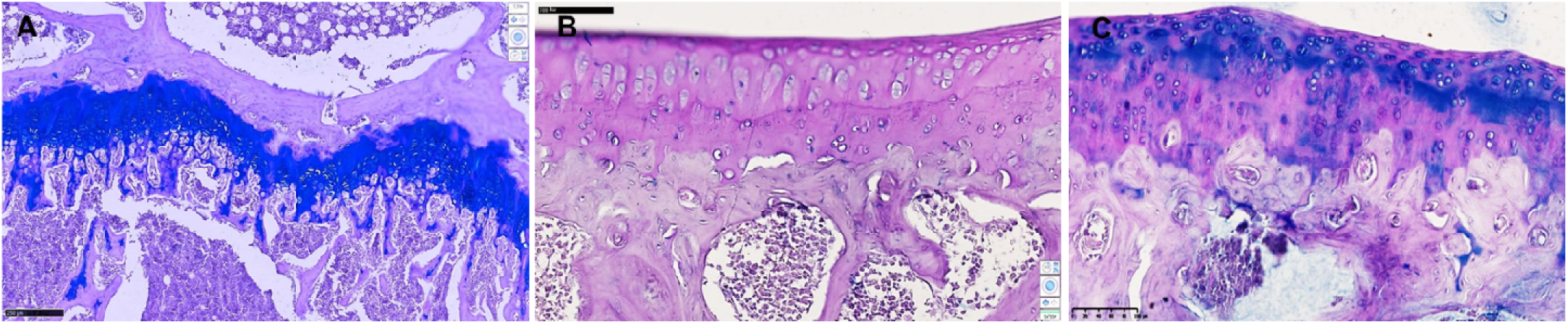
The groups receiving SCM: **A** – Group II (healthy animals and SCM). Isogenic chondrocytes on the articular surface and epimetaphyseal plate form SCHIFF-positive structures of uniform appearance. Schiff-positive structures are found in small quantities in the meniscus and patella. The bone beams in the metaphyseal zone have the same thickness, osteoblasts have a normal appearance, the sizes of the cavities of the spongy substance have the same width. No significant changes were found in other components of the knee joint. **B** – Group IV (trypsin induced knee OA with Estrogen receptor blockade and SCM). Necrobiosis and necrotic chondrocytes are detected in small quantities only. Fibroblast proliferation and synthesis of coarse fibrous collagen are detected only in the subchondral and epimetaphyseal branches. Along the perimeter of the joint, in the pockets of the inner surface of the capsule, disorganization and destructive changes are not detected. On the inner surface of the synovial membrane of the joint, focal hyperplasia of synoviocytes and an increase in regeneration indicators are determined. The joint space is normal in appearance, not damaged, the menisci have the same homogeneous consistency, pathological changes in the structures of hyaline and collagen fibers are not detected. **C** – Group VI (trypsin induced knee OA and SCM). PAS-stained micrographs show focal accumulations of acid mucopolysaccharides, predominantly around the perimeter of chondrocytes and fibroblasts, with pale blue PAS-positive structures. Acid mucopolysaccharides are produced by hyaline chondrocytes and fibroblasts and occur in various forms. The proliferative changes of varying degrees were observed in chondrocytes, as well as the formation of foci of osteofibrosis and calcification instead of foci of osteonecrosis and osteoporosis.

The quantitative histological assessment was scored from 0 to 3, with a total score ranging from 0 (healthy tissue) to 12 (maximum cartilage degradation), summarized in Table 2.

**Table 2.** The severity of cartilage degeneration in different groups according to the modified Mankin scale with a range of scores from 0 (normal) to 3 (severe degeneration).

| MODIFIED MANKIN SCALE |  |  |  |  |  |  |  |
| --- | --- | --- | --- | --- | --- | --- | --- |
| No | Histological parameter | Group I | Group II | Group III | Group IV | Group V | Group VI |
| 1 | **surface structure** (1 point – unevenness, erosion; 2 points – cracks; 3 points – delamination) | 0 | 0 | 3 | 1 | 2 | 2 |
| 2 | **cellular composition** (1 point - slight decrease in the number of chondrocytes; 2 points - significant decrease in the number of chondrocytes; 3 points - no cells) | 0 | 0 | 3 | 1 | 1 | 1 |
| 3 | **staining** (1 point - slight decrease in staining; 2 points - significant decrease in staining; 3 points - no staining) | 0 | 0 | 3 | 0 | 1 | 0 |
| 4 | **cell proliferation** (1 point - isogenic groups of chondrocytes with 2 cells per group; 2 points - isogenic groups with 2-3 cells per group; 3 points - foci of proliferation (more than 3 cells per group). | 1 | 2 | 3 | 2 | 3 | 3 |
|  | **Total:** | **1** | **2** | **12** | **4** | **7** | **6** |

In our study, the lowest score (1) was observed in the control Group I, while the highest (12) was recorded in the untreated Group III (OA + postmenopause). The cartilage condition in Group II was slightly worse than in Group I, while Group IV showed significant improvement over Group III, and Group VI exhibited better results than Group V. A comparative analysis of SCM-treated groups—II (healthy + SCM), IV (OA + postmenopause + SCM), and VI (OA + SCM)—revealed a ratio of 2:4:6, suggesting an ambiguous role of estrogen receptor blockade in postmenopausal OA. These findings cautiously suggest that SCM may exert a more pronounced effect in a postmenopausal context.

## Discussion

The study of on epigenetic reprogramming using small chemical molecules (SCMs) for the rejuvenation of cell cultures (Yang JH et al., 2023) inspired us to conduct this research. The primary goal was to experimentally confirm the effectiveness of SCMs at the whole-organism level and to comprehensively assess their impact. Experimental translational models used in preclinical studies serve as a crucial filter, ensuring that only safe and effective therapies progress to clinical practice.

Aging female rats were selected as the translational animal model, and osteoarthritis (OA) was chosen as the pathological condition. This choice was based on the fact that the aging process in rats is accelerated, and spontaneous OA development in these animals is extremely rare. Therefore, induced models of age-related OA allow for obtaining reliable results in a short time and at reasonable costs. Only rats that met strict selection criteria were included in the study: age over 2 years (equivalent to over 60 years in humans) and weight between 250–270 g (indicative of grade 1 obesity, a known risk factor for OA in humans). The experiment lasted 60 days, corresponding to approximately 6–7 years of clinical observation in human studies.

To investigate the influence of hormonal status on OA progression, the estrogen receptor inhibitor clomiphene was used. Clomiphene competitively binds to estrogen receptors in cartilage tissue (Kalashnikova S.A. & Novochadov V.V., 2009), effectively modeling the postmenopausal state.

### Selection of an Experimental OA Model

To reproduce the pathological process as closely as possible to human age-related osteoarthritis, various OA modeling methods in animals were considered (Kalbhen DA., 1987; Kim M, Rubab A, Chan WCW & Chan D., 2023; Moody HR, Heard BJ, Frank CB, Shrive NG & Oloyede AO., 2012; Salo PT, Hogervorst T, Seerattan R, Rucker D & Bray RC., 2002; Yeh T et al., 2008). These include:

1. **Mechanical damage** (e.g., ligament rupture, meniscus removal)
2. **Spontaneous OA development** (genetically predisposed models)
3. **Intra-articular administration of chemicals**

Chemical induction of OA was deemed the most suitable method, as it allows for the replication of the chronic disease progression characteristic of human OA, closely resembling stage III osteoarthritis according to clinical classification (Korochina KV, Chernysheva TV, Polyakova VS & Korochina IE., 2020).

### Effect of SCMs on Chondrocytes

An indirect marker of replicative aging in chondrocytes is the thickness of the tibial epiphysis. With age, this layer thins, triggering a cascade of pathological changes characteristic of OA. Interestingly, in all groups receiving SCMs, the thickness of the epiphyseal cartilage was significantly greater than in the intact group without OA (Fig. 14). Moreover, histochemical analysis revealed a sharp increase in mucopolysaccharide synthesis in these groups, confirming not only the restoration of damaged joints but also an increase in the metabolic activity of aging chondrocytes.

Another criterion for assessing chondrocyte maturity is the N/C ratio which was calculated in mitotically active areas of cartilage (see Fig. 15). Data analysis demonstrated a significant change in this indicator under the influence of SCMs in both the superficial and intermediate zones of the articular cartilage. This suggests epigenetic rejuvenation of chondrocytes in the most vulnerable areas affected by OA.

Importantly, no quantitative increase in the number of chondrocytes was observed. However, their metabolic activity was notably enhanced, as evidenced by increased cell vacuolization and intensified staining of the intercellular matrix with Schiff reagent. This indicates activation of extracellular component production. Such findings suggest that the SCMs used in this study do not exhibit oncogenic potential, a crucial safety criterion.

#### Unexpected Results

One of the most unexpected findings was the differing responses to therapy between the groups receiving SCMs with estrogen receptor blockade and those without. According to the Mankin scale, the group of rats with OA and postmenopause that received SCMs demonstrated the best outcomes among all the groups with induced OA. However, the histological analysis showed that untreated OA in the groups with estrogen receptor blockade resulted in the most pronounced degenerative changes.

These results suggest that estrogens may play a dual role in regeneration: while their deficiency exacerbates degenerative processes, their presence might also limit the rejuvenating effect of SCMs. This hypothesis warrants further investigation, as elucidating the mechanisms underlying the interaction between estrogens and epigenetic reprogramming could significantly influence therapeutic strategies for treating OA in postmenopausal women.

Thus, our findings confirm the potential of epigenetic reprogramming in the treatment of age-related OA and open new avenues for exploring the interplay between hormonal regulation and regenerative processes.

## Conclusions

1. A model of induced osteoarthritis in rats was developed and validated, enabling rapid and effective assessment of epigenetic reprogramming under the influence of SCMs in vivo.
2. Epigenetic reprogramming of chondrocytes using the selected SCMs demonstrated high efficacy in the induced OA model in aging rats.
3. Histological analysis revealed cartilage tissue regeneration, increased epiphyseal layer thickness, and enhanced mucopolysaccharide synthesis in the groups receiving SCMs.
4. The rejuvenating effect was accompanied by activation of metabolic processes in chondrocytes, as evidenced by changes in the N/C ratio and increased extracellular matrix production.
5. Estrogen receptor blockade altered the response to SCMs, highlighting the need for further investigation into the potential interactions between estrogens and epigenetic rejuvenation mechanisms.

## Materials and Methods

### Laboratory Animals

Experiments were conducted on female white rats (*Rattus norvegicus albinus*) weighing 250–270 g. The selected rats were fertile females, classified as at least class 2 (having birthed no fewer than five pups per litter), with estrous cycles occurring every 6–10 days. The animals were housed and fed under standard vivarium conditions throughout the study, adhering to the recommendations of the Ethical Committee of Uzbekistan (2014) and the Ethical Guidelines for the Use of Animals in Research (2019). The Ethics Committee’s permission to work with laboratory animals is enclosed.

### Small Chemical Molecule Set (SCM)

SCMs were administered at concentrations (Table 3) previously established in in vitro VC6TF experiments for epigenetic reprogramming of cells (Yang JH et al., 2023).

**Table 3.**
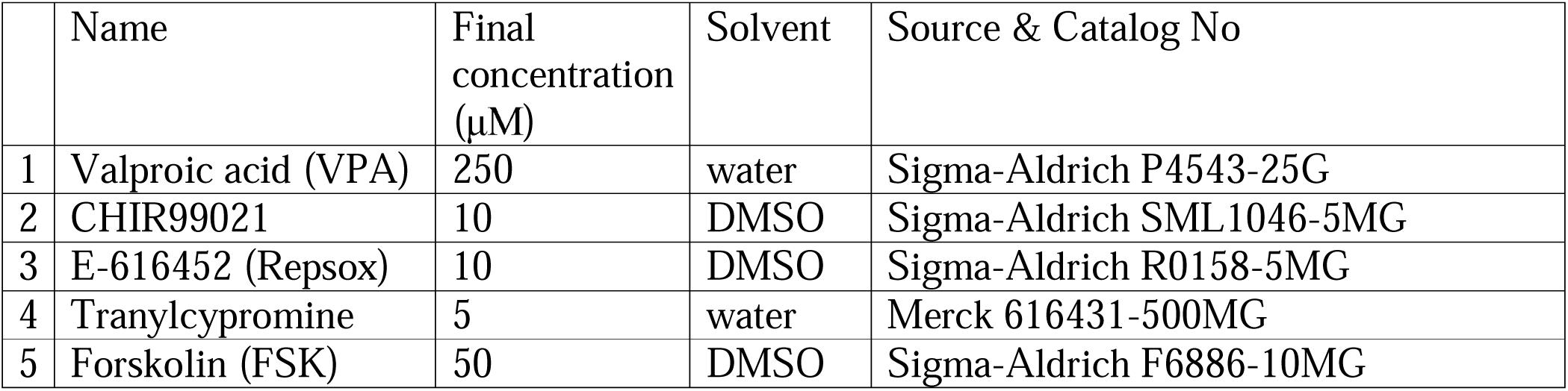
Composition and concentrations of small chemical molecule (SCM) cocktail.

### Modeling of the Postmenopausal State

To simulate the postmenopausal state, clomiphene citrate, an estrogen receptor blocker, was used. As previously demonstrated (Kalashnikova S.A. & Novochadov V.V., 2009), this blocker competitively binds to chondrocyte estrogen receptors, effectively inducing menopause in experimental rats.

### Histological and Histochemical Methods

Following euthanasia, the right knee joints, along with the lower third of the femur and fragments of the tibia, were dissected for histomorphological examination of cartilage and bone tissue (Muzhikyan A.A., Shekunova E.V., Kashkin V.A., Makarova M.N. & Makarova V.G., 2018; Perepelkina D.O, Nasibova S.S, Bakhtin V.M. & Izmozherova N.V., 2019). Three histological and histochemical staining methods were employed to evaluate potential epigenetic reprogramming of chondrocytes: hematoxylin and eosin (H&E), Van Gieson staining, and the periodic acid-Schiff (PAS) reaction. These methods selectively highlight key structures essential for analysis.

The collected biomaterial (right lower limbs) was coded and stored in separate containers according to experimental groups. Tissue processing followed standard paraffin embedding protocols using Histomix paraffin (BioVitrum, Russia), and 5–8 μm sections were prepared with a microtome.

Microscopic analysis was conducted using a ZEISS Primo Star light microscope, and microphotographs were captured. An independent histopathologist, blinded to the experimental groups, documented digital images of histological slides. Slides were analyzed using a NanoZoomer Hamamatsu light microscope (Japan) at 50x, 100x, 200x, and 400x magnifications. Digital imaging was performed using a NanoZoomer digital camera and software (REF C13140-21.S/N000198/HAMAMATSU PHOTONICS/431-3196 JAPAN).

A morphometric study of tissue structural units was conducted using a multiplex confocal morphometric device. The QuPath-0.5.0 software was used to generate two-dimensional images, which were analyzed along the X and Y axes to determine tissue area and lesion location. Articular cartilage zones were marked to assess joint tissue involvement in the pathological process.

### Severity Assessment Using the Modified Mankin Scale

Cartilage degeneration severity was quantified using the modified Mankin scale (Moody HR, Heard BJ, Frank CB, Shrive NG & Oloyede AO., 2012), with scores ranging from 0 (normal) to 3 (severe degeneration). The following parameters were analyzed:

- **Surface structure:** (1) unevenness/erosion, (2) cracks, (3) delamination.
- **Cellular composition:** (1) slight decrease in chondrocytes, (2) significant decrease, (3) absence of chondrocytes.
- **Staining intensity:** (1) slight decrease, (2) significant decrease, (3) absence of staining.
- **Cell proliferation:** (1) isogenic chondrocyte groups with two cells per group, (2) groups with two to three cells, (3) proliferation foci (>3 cells per group).

A maximum pathology score of 12 indicated the most severe degenerative changes.

### Statistical Analysis

All data were subjected to statistical and mathematical analysis. Quantitative indicators were used to assess reliability, with results expressed as corresponding significance levels. Statistical processing was conducted using MS Office Excel 2007 and STATISTICA (Windows 10) software, following standard statistical analysis methods.

Dynamic changes within groups and across the population were assessed using **Student’s t-test**. Pearson’s correlation coefficient was calculated for normally distributed data, while Spearman’s correlation coefficient was used for non-normally distributed data.

